# *Thinging* Through Modelling. Active Inference meets Material Engagement

**DOI:** 10.1101/2025.08.17.670756

**Authors:** Laura Desirèe Di Paolo, Bruno Vindrola-Padrós, Andy Clark, Axel Constant

## Abstract

In this simulation study, we adopt the comprehensive neurocomputational approach of Active Inference (AIF) to illustrate some key concepts of Material Engagement Theory (MET) [1]. MET posits that craftwork does not require, or rely on, rich internal ‘pre-planning’, i.e., complex and highly detailed representations that occur mainly in the maker’s head. Instead, the maker engages materiality through *’thinging’*, where the human agent (the maker) is guided by and leverages the materiality of the artefact (such as a spinning of clay or a chunk of marble). MET assigns a crucial co-participatory role to materiality, attributing agency to it. We investigate MET’s claims through the widely adopted theory of AIF [2]. Our first aim is to simulate the plausibility of the (creative) *thinging*, adopting a simple modelling scenario. Then, we also discuss its applicability to other, more complex cases, which seem to require greater levels of ‘pre-thinking’ (e.g., planning, imagining and conceptualising outcomes). We highlight how these cases, too, align with the general principle of *thinging*. With our AIF neurocomputational understanding, we explain that even in these situations, the predictive brains involved in the creative process attempt to minimise the complexity of their internal model. The upshot of this is that, always and everywhere, our human minds engage the materiality to make the most of the characteristic dynamics of the world surrounding us - things and processes alike.

## 1 INTRODUCTION

### 1.1 This paper

This paper has two main aims. First, it presents a “proof of concept” simulation study utilising Active Inference (AIF) -- a computational approach in theoretical neurobiology that implements the theory of predictive processing and the predictive mind [3,4] -- to illustrate key concepts of Material Engagement Theory (MET) in the context of craftwork and artefact-making [1]. Second, the paper argues that the computational tools used in our simulation can be used to bring support to some of MET’s claims. The AIF simulation that we present illustrates two of MET’s central claims: that, (i) artefact-making activities like pottery seldom (if ever) require rich “pre-planning”, in the form of complex and highly detailed mental representations that occur solely in the maker’s head, prior to, and independent of, any engagement with the material. Instead, during crafting activities, (ii) makers leverage and are guided by the intrinsic dynamics entrenched in the materiality of the artefact itself (e.g., in throwing a lump of clay) [5]^1^. These claims, at the core of MET, understand ‘thinking’ processes as *thinging*, and attribute a driving role to materiality itself in creative activities (i.e., ‘creative *thinging*’) [12].

We rely on AIF because it has gained attention as a plausible model of the human brain, while attributing a fundamental role to mind-body-environment interactions. Specifically, it is a theory in computational neurobiology that views cognitive functions such as perception, attention, learning, and action as processes of embodied inference [2,13]. The AIF has been implemented in computational modelling methods that enable the simulation of many forms of simple and goal-directed behaviour (for a review of the empirical evidence, see: [14]). Recently, the application of AIF has been extended to the modelling of dynamics unfolding beyond the skull, including social dynamics [15,16], evolutionary dynamics [17,18], and, importantly for our purpose here, human-material interactions [19–23]. AIF also offers a powerful framework for thinking about how intelligent agents spot and exploit affordance structures in the local environment: the opportunities that the world presents for structured interactions that move us closer to our goals ([24–26]).

### 1.2 The simulation

Our simulation illustrates MET’s thesis of creative *thinging* employing a simple numerical simulation, where the agent does not have access to higher-level cognitive skills for sophisticated pre-planning. The task of the simulated human agent was to perform a series of actions that would yield an apt and maximally aesthetic vase by carving the lump of clay up or down in different locations, based on the action that it perceives at that moment in time as being afforded by the lump of clay. In line with MET, we aim to depict successful pottery manufacture as heavily relying upon the features of the material environment, which ultimately affords the kind of actions leading to the execution and finalisation of pottery making. AIF and MET find common ground in the concept of affordances (i.e., action possibilities offered by the material and perceived by the human agent), yet they work complementarily. While MET emphasises the interaction between the artefact and environment affecting the agent, AIF delineates affordances as attributes of actions, highlighting the human agent’s expectations and capacity to navigate uncertainties presented by the material [26,27]. Eventually, complex culturally acquired behaviours are triggered by these affordances [28,29]. We explore the application of *thinging* to these situations that supposedly might require extensive pre-planning and detailed mental visualisation of desired outcomes. In line with MET and based on our numerical simulation, we show that, no matter how complex the final product might look, the *thinging* process is always at work. The reason, we argue, is to be found in the very nature of the human mind and the predictive brain.

The analysis of crafting activities must retain the cultural context in which they occur and develop. Similarly, in our work, the simulated potter approaches the clay with some broad ‘cultural criteria’, reflecting the culturally informed notion of what they are setting out to produce. To achieve these cultural criteria, the potter works the clay using a specific forming technique to create a vase that must meet specific quality standards (i.e., *aptness*) and maximise a broadly determined aesthetic. Even if in broad and simplified design requirements, the cultural criteria reflect the cultural sensitivity to certain shapes or processes, influencing attention and resulting in cultural biases and affordances (see, e.g., [15]^2^. The aptness criterion measures how effectively a product solves the problem it was designed to target. In our simulation, the aptness criterion refers to design requirements that the vase must possess, namely, the ability to stand and function as a container, together with having a symmetrical shape and a conical neck [34]. Choosing among many possible design classifications, we adopt Birkhoff’s measure of aesthetic value, which is determined by the complexity of “vase-like” artefacts upon the number of breaking points, or points of changes in directions along the contour line of their silhouette [35–37]. Again, for the sake of the argument, our simulation assumes that the increase in the number of symmetrical features is considered of greater aesthetic value within our population of potters (i.e., the cultural standards of such a population). Lastly, our simulation considered that the potter utilises a culturally defined technique to shape the vessel, traditionally termed “paddle and anvil”. This technique, widely adopted in handmade pottery, involves placing an anvil - usually a flat or gently curved surface, such as a wooden or stone tool - in the interior of the lump of clay, and hitting or pressing it from the exterior with a ribbed or smooth paddle (see [38].

Once these broad aims were installed, our first simulated pot maker had a set number of trials to complete the task corresponding to the minimal number of trials required to produce a maximally apt and aesthetic vase (see: Method, Sec. 2). We ran 6 simulation series and manipulated two parameters: (i) Attention level, or precision of the potters’ model, and (ii) the stability of the potting material implemented as the randomness of the observations stream, here representing affordances or action possibilities. Our results show that greater aptness and aesthetic value can be achieved *only* with the cooperation of material agencies. This is particularly evident in cases where the material behaves unexpectedly (e.g., random observation) and when complex plans are not available, such as if a potter works with types of natural clay that contain unwanted inclusions, unpredictable shrinkage, and unreliable thermal properties. That is, the production of suitable vases (*aptness*) depends significantly on the material’s ability to generate suitable affordances that support and encourage pot-making activities. However, as we show in the Results (Sec. 3) and in the Discussion (Sec. 4), and as every artist, artisan, and archaeologist surely knows, sometimes *just enough* unpredictability and “uncooperativeness” of the material (compared to when the material behaves passively, indicating exactly how it ought to be engaged) enables the creation of artefacts with a greater aesthetic value, as also shown by our potting agents. Section 5 considers different cases, including those where the potter has both pre-defined goals and the necessary expertise to realise them. We will argue that even in those situations, *thinging* dynamics are essential to obtain a successful outcome. Additionally, we discuss the limitations of applying a formal framework such as the AIF to replicate (some) *thinging* dynamics, that could never be fully formalised.

## 2 METHOD

### 2.1 The task

We simulate a pot-making activity using a heuristic that involves the interaction between an AIF agent and a ‘material’ agent (a lump of clay). We depict how this interaction might indeed produce sophisticated forms of artefacts, in the absence of complex pre-planning “in the head” (e.g., with deep hierarchical models). The model for the human agent is an industry standard generative model under AIF, implementing a Partially Observable Markov Decision Process (POMDP), with action policies allowing potting agents to work different locations of a lump of clay that can be hit or pressed, and decorated [39,40]. The material agent (i.e., the clay) is simulated only implicitly, as that which delivers a certain stream of sensory stimulation to an AIF agent. In this way, while those stimulations are contingent upon the actions of the potter, they also reflect properties and proclivity of the material, i.e., the agency of the clay. We thus model the material agent as a generative process: a description of the dynamics of the world generating sensory observations in response to how an agent moves in the world and acts on it. This is a useful simplification that is arguably conceptually well-motivated. Indeed, since a potter can only interact with the material order through an exchange of sensory observations, whatever happens to the potter agent in virtue of the world’s materiality must be conveyed through the sensory stream of the agent. This licenses us to treat materiality, under AIF, as a sensory stream produced by a generative process. It might be appreciated then, that the potting agent could equally be reduced to a stream of, this time, “actions” which can be viewed as the “sensations” of the material environment (e.g., lump of clay) [16]. Eventually, our simulation boils down to an interaction between a potter and a lump of clay mutually shaping each other through bidirectional dynamics and sensory-motor couplings.

We simulated 100 such agents under 4 conditions (see section 2.2), and the simulated potter had 12 trials to complete the task of forming an apt and aesthetically pleasing vase. The models being deterministic, and the only element of stochasticity contained in our simulation being the randomisation of observations input to the model under certain conditions, 100 agents per condition were enough to arrive at results that could illustrate the dynamics that we were interested in and that are characteristic of AIF agents. We predefined the aptness and degrees of aesthetic of the pots that our agent could achieve (figure 1) based on the set of prior beliefs and likelihood implemented by the model (figure 2). Pots number 2, 5, 8, and 11, such as depicted in figure 1, were considered apts. The most complex one (i.e., number 11) was considered the most aesthetically valuable. The possible actions afforded by the lump of clay - i.e., observations - were the pressing or hitting ‘downwards’, ‘upwards’, or ‘decorating’ of a section. Narratively, the sequence of events implemented by the for loop of our simulation was:

i. the perception of an affordance (i.e., observation), which indicated which of the three actions was available (e.g., “I see that the clay in its current form affords decorating, given the 12 locations on the lump of clay where decorations could be applied”),
ii. the inference of the future state, or location on the piece of clay where such an action should be performed (e.g., “I think that the decoration should be located in the centre of the piece of clay”); and
iii. the prediction of the next affordance and the enaction of the prediction (e.g., “I predict that decorating is what comes next, and therefore I will decorate the location that I inferred”) (see Figure 1)^3^. Note that given the simplicity of our model, the notions of “inference” and “prediction” remain rather minimal. As depicted in Figure 2, they are predictions of an action possibility based on the inference of an action to be performed. In that sense, the inference utilises representations of actions in order to predict action, not at all describing content to direct contentful plans. They involve representations akin to what Ruth Millikan called Pushmi-Pullyu, “serving at once to direct action and to describe one’s future so that one can [act on, rather than plan around,] it” [41].

**Figure 1.**
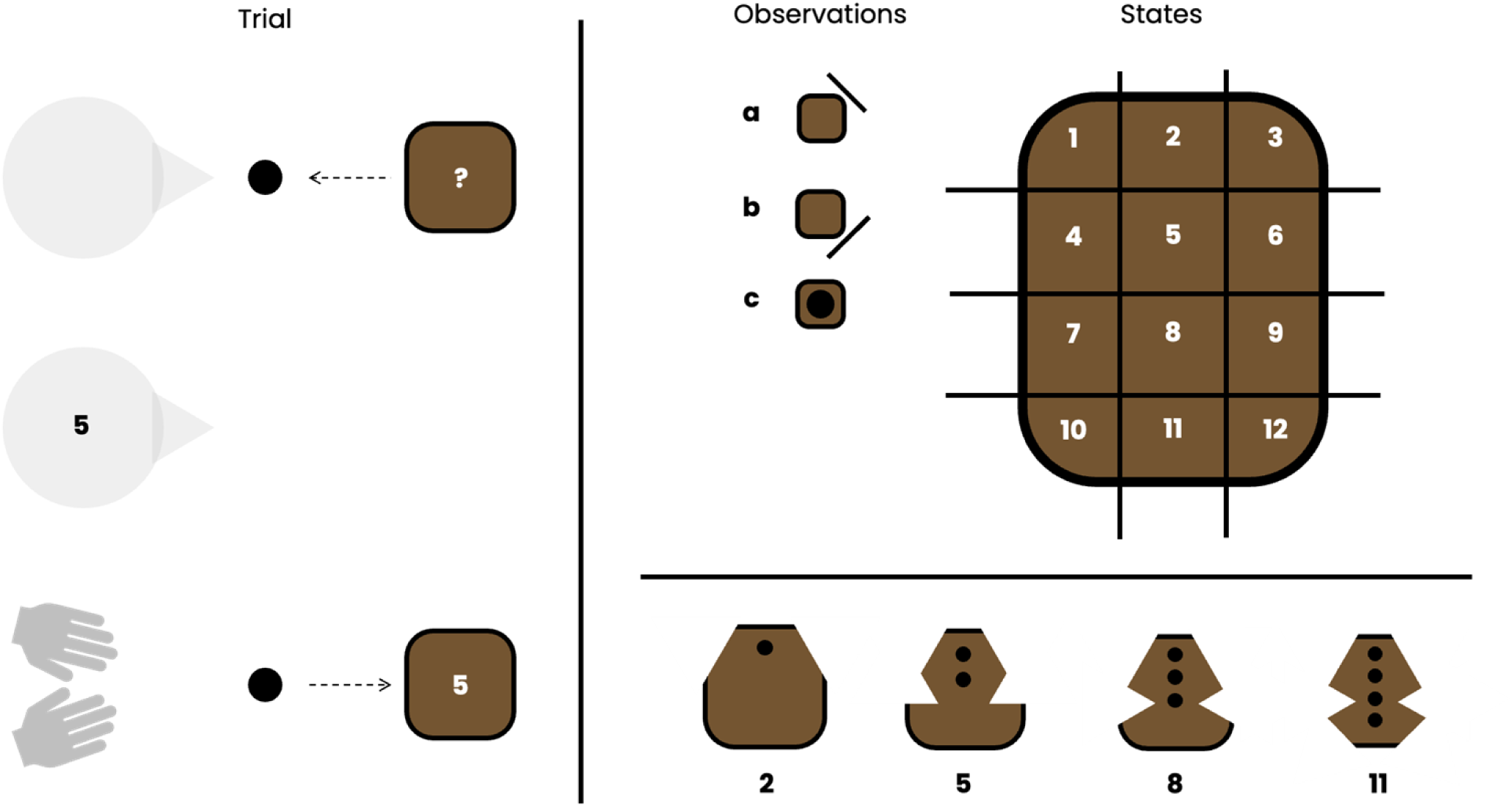
**Top right panel:** The generative model of the agent represents the structure of the world in terms of “states-to-be-acted-upon” and observations, or affordances. The states of the world correspond to the 12 possible locations of the lump of clay that can be acted upon. The observations correspond to the 3 possible affordances that can be generated by the lump of clay, namely (a) “down pressability”, (b) “up pressability”, and (c) “decoratability”. Each location affords a different action (i.e., an “ability). **Bottom right panel:** over the 12 trials, the agent can produce a variety of different shapes, 4 of which are considered apt (pots 2,5,8 and 11). The 4 apt pots are ranked in terms of their aesthetic value, broadly construed (cf. Birkhoff aesthetic value). Pot 2 is the simplest, thus the least aesthetically valuable, and Pot 11 is the most complex, most aesthetically valuable pot. **Left panel**: from top to bottom, each trial has three moments (i) the perception of an affordance, (ii) the inference over which of the location may have produced the affordance, and (iii) the prediction of the next affordance (i.e., observation), and associated location (i.e., state); the predicted affordance determining the nature of the action, and the predicted state corresponding to where the action will be. This means that, contrary to orthodox modelling approaches in active inference, we consider the “observation” as the source of action, instead of the sequence of states (e.g., moving from pot location 1 to 2).

**Figure 2.**
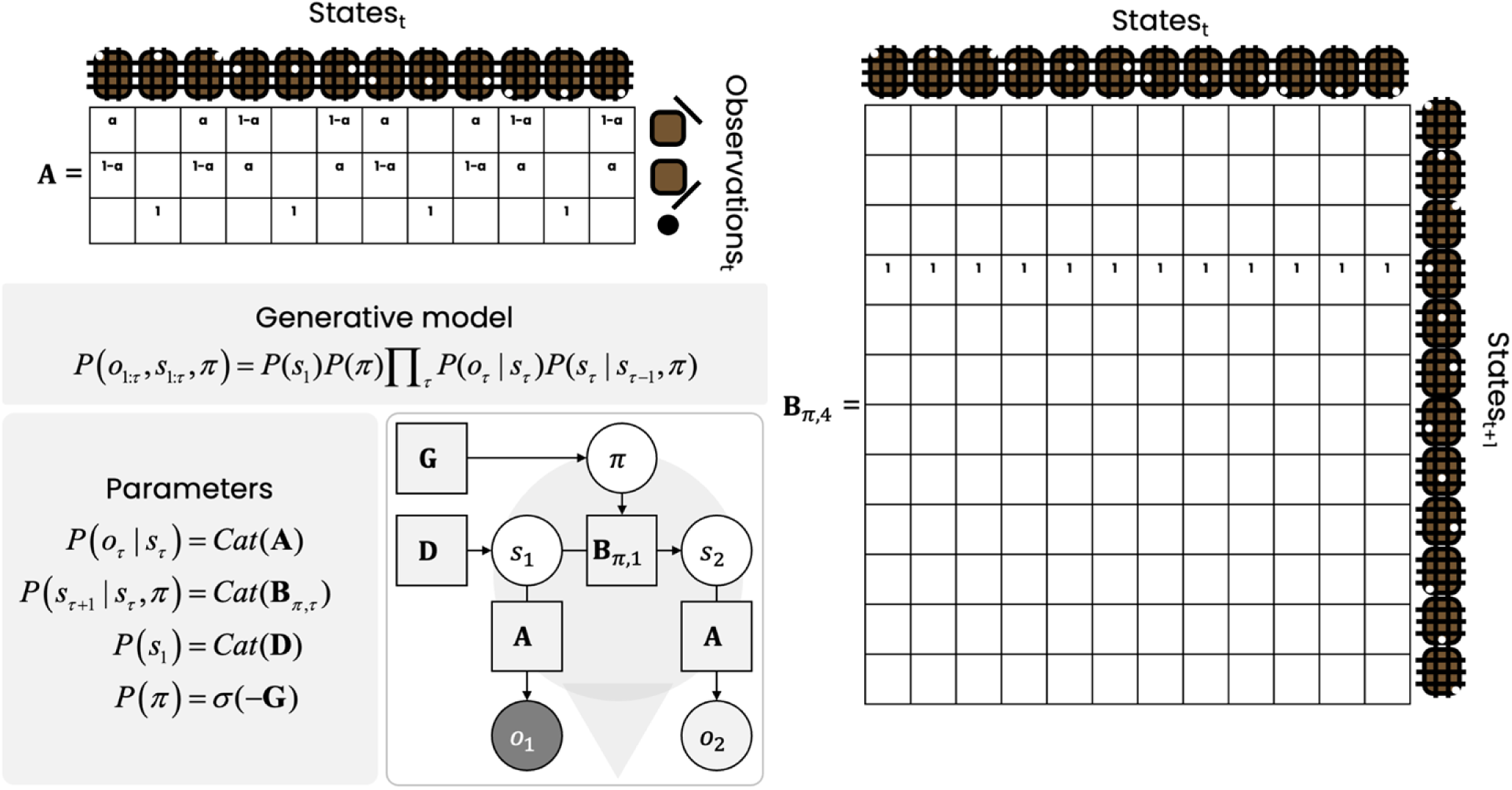
**Bottom left panel:** The generative model of the potting agent decomposes into parameters that correspond to priors B, D and G, and a likelihood A. White circles represent random variables such as the hidden states and the policy to be inferred. The hidden states correspond to the 12 possible locations on the pot, and the policies have a 2-time-step depth and can go from the current location to the next location out of the 12 possible locations. Grey circles represent the outcomes, or observations, which in our case refer to the affordances of the pot (downward-capability, upward-cuttability, or decorability). In each trial, an outcome is presented to the agent, and the next outcome has to be inferred along with the policy that will lead to the state thought to afford that outcome; hence, the outcome o2 is represented as a white circle. Squares represent the model parameters A, B, D and G. The parameters are categorical densities (*Cat*). The policy (pi) that is inferred is the one that minimises expected free energy (G), which includes a parameter “C” that indicates which is the preferred outcome for each trial. For a description of the update equations and underlying theory, see: [40,49]. In our simulation, the agent always had a preference for the outcome that corresponded to an affordance (i.e., observation) that matched the ideal pot (e.g., having a preference for downwards cuts at locations 4, 6, 10 and 12). **Top left panel**: The likelihood parameter A maps the three possible outcomes onto the 12 possible location states. Under conditions C2, C3, and C4, we gradually lowered the precision. This was achieved by changing the precision of the mapping between states at the edge of the lump of clays (i.e., 1,3,4,6,7,9,10,12) from a = .9 (high) to a = .8 (mid) and to a = .7 (low). We stopped at 7 because, as of a = .6, none of the agents could complete any of the 4 possible pots anymore. **Right panel**: The agent used 12 different policy-dependent state transition matrices (B), each allowed for the enactment of one of the 12 possible policies. The one depicted in Figure 2 corresponds to policy “4”, as it attributes 100% probability of moving from each of the possible 12 states at time “t” (columns) to state 4 at time “t+1” (rows).

### 2.2 Conditions

The simulation was run under 4 conditions, each of which manipulated one or both of the interesting parameters for this study. The first parameter corresponded to the ‘attention’, or level of precision, attributed by the potter to the observations that functioned as affordances of the clay. Precision refers to the uncertainty in the mappings between observations and a state of the likelihood. In AIF, exogenous attention is modelled as the susceptibility of updating Bayesian beliefs about the conditional relation between an observation and its causes, or state, and depends on the precision, or inverse covariance, of that mapping [42]. According to AIF, and known as ‘precision weighing’ [43], this way of weighing the evidence may be realised by the synaptic modulation of neurotransmitters such as dopamine and acetylcholine in the visual cortex [44]. In the context of this simulation, manipulating attention or precision allows us to model two key cases (and all points in between): in the first, the potter is strongly guided by their own outcome-expectations; in the second, the connection between outcomes and expectations hold more loosely (i.e., less precision), permitting greater agency to the clay itself. Under AIF, these two cases are equivalent to allowing the potter to pay more or less attention to certain representations of actions that she might expect, making her rely more or less on the action afforded by the materiality. That is, high precision turns into making the potter rely more on representations of her own intended actions, and less on the affordances provided by the clay. The second parameter that we manipulated was the randomness of the affordances generated by the clay. To achieve an apt and maximally aesthetic vase, the potter needed to respond to the affordances of the clay, which themselves had to appear in an ordered and predictable way, as per the normal sequence of forming the pot by hitting or pressing the clay (e.g., starting at the top of the vase and shaping the edges moving downward). Increased randomness of the observations, or affordances generated by the clay, reflects the unpredictability of certain materials that may be more difficult to work with, as they may not afford expected action during pot-making activities.

Condition 0 (**C0**) functioned as the baseline simulation, with perfect precision and no randomisation of the affordances generated by the pot. Simulation 1 (**C1**) shows what happens when we gradually decrease the potter’s sensitivity to the repertoire of the complex action patterns afforded by the clay (i.e., “affordance uptake”), such as when working with material that is highly unpredictable or offering highly unexpected affordances. The unpredictability was implemented as the randomisation of the observations, which function as input to the model in each of the 12 trials -- the observations corresponding to the affordances as depicted in Figure 1 (i.e., hit down or up, decorate). Simulation 2 (**C2**) shows a situation in which the same decrease in precision is combined with more predictable material (i.e., “semi-unexpected affordances”). The increase in predictability comes from the fact that we randomised the observation only every other trial. Simulation 3 (**C3**) shows how the decrease in precision influences the ability of the potter to reach one of the four apt pots when working with stable material, which affords the kind of action expected for each location of the lump of clay. Overall, the simulations suggest that AIF is a highly flexible modelling tool capable of capturing arbitrary degrees and the combinations of both sides of the human agent’s knowledge and skill, and the material agent’s properties and affordances.

### 2.3 The generative model

The model of the potters was a generative model specified as a joint probability (P(S,O)) of world states “S” (S_t_ = (s_1_,s_2_,…s_t_)) and of sensory observations “O” (O_t_ = (o_1_,o_2_,…o_t_). The states corresponded to the 12 possible locations on the clay, and the observations corresponded to its affordances: “upward-pressability”, “downward-pressability”, and “decorability”. Over the 12 time steps, the combination of inferred states and observations could yield 4 possible completed pots (Figure 1). Prior beliefs over initial states (denoted as P(s1), or D) and prior beliefs over -- policy dependent -- state transitions for each possible policy ‘pi’ (denoted as P(S_t+1_ | S_t_,pi), or B) are combined with the likelihood mapping between states and observations (P(O|S), or A) to infer the current state upon the receipt of an observation, and predict the future state and the associated observation. The variational message passing algorithm that applies to our partially observed Markov decision process (POMDP) and that allows inference is described in [40].

The inference of the states (here: the locations of the lump of clay worked by the potter) is done through minimizing variational free energy (i.e., maximizing Bayesian model evidence [45]), which effectively minimises the discrepancy between predicted observations (i.e., the affordance expected by the potter) and current observation (i.e., the current affordance). The prediction of future states is achieved by selecting the policy with the greatest model evidence expected under the future states at t+1, such as encoded by the policy-dependent transition matrix. This means that for each policy, there exists a transition matrix. In this simulation, the potter had access to 12 transition matrices, each corresponding to the transition from any of the 12 possible locations to one of the 12 possible locations of the lump of clay. The prediction of the future state thus simply amounts to selecting the policy that has the least expected free energy G_π_, or free energy expected under the posterior beliefs about observations for an action. Expected free energy is a mixture of expected information gain (cf. the quantity used in active learning and Bayes optimal experimental design) [46,47] and expected value (cf. the quantity used in optimal Bayesian decision making) [48].

In our simulation, the action corresponding to the simulated potters’ action (i.e., working the clay at one of the 12 possible locations) was the expected free energy minimal action. Under each condition, each of the 100 simulations ran separately followed the following steps: (i) Setting the precision over the likelihood matrix to the desired level, depending on the condition at hand, (ii) iterating using a “for loop” over the 100 agents, 12 trials per agent, by randomizing the observations under condition 2, 4 and 6 for each of the 12 trials, and passing the parameters such as described in figure 2 to the MATLAB routine spm_MDP_VB_X.m. We stored the belief updates of the 100 agents in a structure and performed the statistical analyses presented in the following section on the stored data.

## 3. RESULTS

As defined above, the task of the potters was to achieve the most apt and aesthetic vases. Four such vases were possible under the circumstances, i.e., pots 2,5,8 and 11 (depicted in Figure 1). We ran 100 agents per condition, each agent having access to 12 trials to complete a maximally apt and aesthetic vase (i.e., vase 11). The condition **C0** was our baseline condition in which affordances were stable, and the precision of the agent was perfect (i.e., a = 1, cf. figure 2). This condition systematically yielded 100% success for all the agents (i.e., the 100 agents reached pot number 11 within the 12 trials). Below we show the between-conditions (Figure 3) and within-conditions results (Table 2). Between-conditions results, look at which of the conditions yielded the most apt and aesthetic pots. Within-conditions results look at the loss rate and the mean rate of change of agents in each condition. The loss rate refers to the percentage of agents who have completed pot 2 but have failed to complete pot 11. The mean rate of change is the mean rate of change of the number of agents having completed pot 2, then pot 5, then pot 8, then pot 11 (i.e., the mean of “pot2-po5/pot5-8” and “pot5-po8/pot8-11”). The mean rate of change reflects the curves of the bar charts in Figure 3.

**Figure 3.**
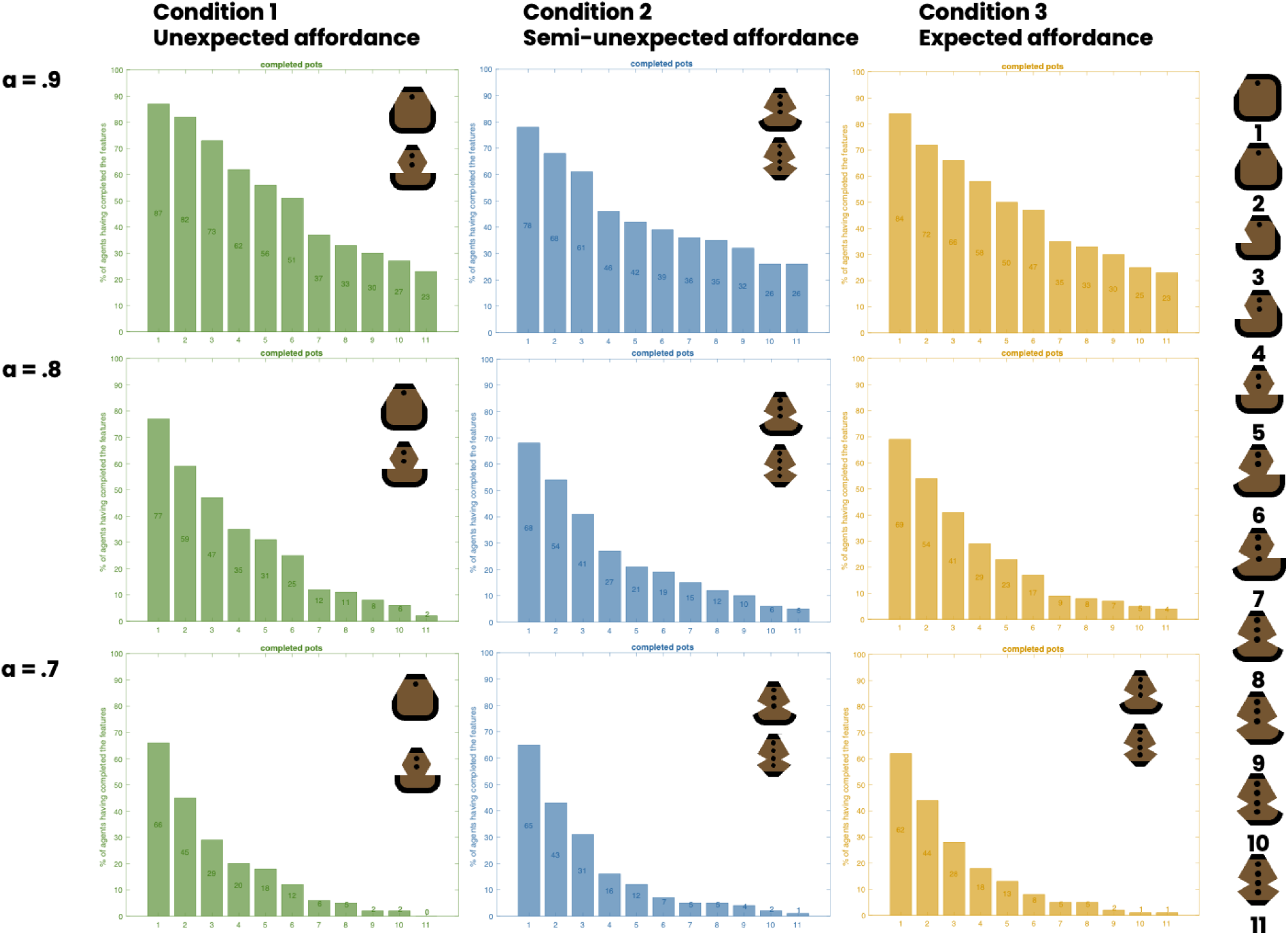
Between-conditions results. For each plot, the X axis represents the pot number, and the Y axis represents the % of agents having achieved the pot corresponding to the pot number.

**Table 2.**
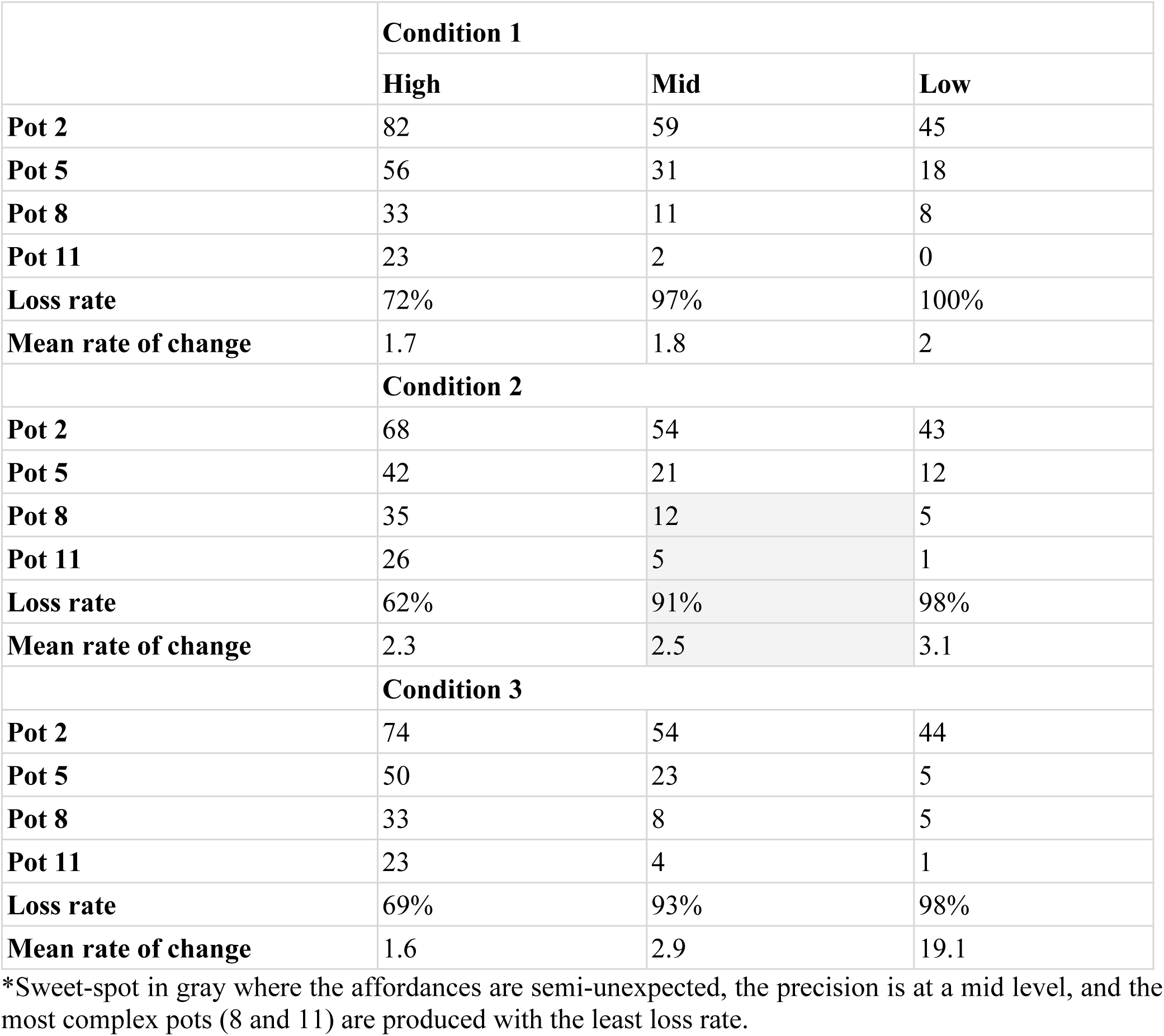
Within conditions results.

|  | **Condition 1** |  |  |
| --- | --- | --- | --- |
|  | **High** | **Mid** | **Low** |
| **Pot 2** | 82 | 59 | 45 |
| **Pot 5** | 56 | 31 | 18 |
| **Pot 8** | 33 | 11 | 8 |
| **Pot 11** | 23 | 2 | 0 |
| **Loss rate** | 72% | 97% | 100% |
| **Mean rate of change** | 1.7 | 1.8 | 2 |
|  | **Condition 2** |  |  |
| **Pot 2** | 68 | 54 | 43 |
| **Pot 5** | 42 | 21 | 12 |
| **Pot 8** | 35 | 12 | 5 |
| **Pot 11** | 26 | 5 | 1 |
| **Loss rate** | 62% | 91% | 98% |
| **Mean rate of change** | 2.3 | 2.5 | 3.1 |
|  | **Condition 3** |  |  |
| **Pot 2** | 74 | 54 | 44 |
| **Pot 5** | 50 | 23 | 5 |
| **Pot 8** | 33 | 8 | 5 |
| **Pot 11** | 23 | 4 | 1 |
| **Loss rate** | 69% | 93% | 98% |
| **Mean rate of change** | 1.6 | 2.9 | 19.1 |

\*Sweet-spot in gray where the affordances are semi-unexpected, the precision is at a mid level, and the most complex pots (8 and 11) are produced with the least loss rate.

The results show a clear trend: as precision decreases, fewer agents reach pot 11 across all conditions. Condition 1 enabled the most agents to complete pots 2 and 5, indicating it was easier for agents to achieve pots considered less apt and aesthetic. This trend was influenced by precision, where decreasing levels narrowed the gap between conditions in terms of the number of agents reaching pots 2 and 5. More complex pots (pots 8 and 11) were most frequently achieved under condition 2, which allowed for some observation instability, though less so than condition 1. Although condition 3 provided consistent affordance with no instability, it produced poorer results than condition 2, except at low precision (a = .7). Overall, among the results within-conditions is found that lower precision results in higher loss rates, though this effect is least pronounced in condition 2 (semi-stable affordances). The mean rate of change from high to low precision increases systematically from condition 1 to condition 3, suggesting that more stable affordances (with non-ideal precision, i.e., less than a = 1) lead to fewer agents transitioning from basic pots (e.g., 2 and 5) to more complex pots (e.g., 8 and 11)^4^.

## 4. DISCUSSION

We now discuss the significance of our results in relation to the notion of cognitive *thinging*, and more broadly, what seems to motivate the intellectual project of MET. MET claims that neural, bodily and environmental factors combine in the making of human artefacts. In this complex interaction, artefacts acquire agency since they are structures with their own complex dynamics that impose constraints on, and drive the creative activity of artefact makers - hence, the notion of *thinging* [1,12,50–52]. In our simulation, the agency of the lump of clay was implemented as the (un)expectedness of the action afforded by the clay, or precision of observations; the clay displaying agency under the form of “resisting being shaped in a way that matches the potter’s prediction”. On the other side of the MET story, we add, the cognitive system of the human maker (the other agent) engages with the material in ways that make the most of the characteristic dynamics of structured interaction spaces (e.g., [21,22,53,54]. In our simulation, this was implemented as a minimal -- shallow -- model endowed with a representation of actions, not a rich description of world properties. MET relies on the concept of ‘metaplasticity’ to explain the ongoing process of developmental and evolutionary ‘incorporation’ of extra-bodily elements within the cognitive system of the human agent, a process defined as ‘Human Becoming’ (see: [51,52,55,56]). We aimed to elucidate this process, which is central to the complex dynamics intertwining human and nonhuman agents in artefact-making (see, e.g., [19,20].

MET is motivated by the problem of artefacts’ ‘Ontological Boundaries’. Archaeological theories consider artefacts simultaneously "actions", "objects", and "traces" [50]; actions because their production is a process (e.g., potting); objects because they are outcomes of this process; traces because they carry in them signs of, and are considered to represent their productive process. The “traditional” view, epistemologically based on a clear separation between brain, body, and environment, establishes a default system of cognitive boundaries in which the three elements are considered isolated components. This epistemological separation becomes ontological when we try to understand which factor has independently contributed to the production of the artefact, and at which point during the process. As if each of them could be related to the outcome *independently* of the co-occurrence of the others. Additionally, the traditional view separates the artefact, the out-of-the-body object, from the mental activity or pre-planning, the in-the-brain activity. Ultimately, the artefact becomes merely the representation of such a mental activity which precedes and causes the materialisation of the object in the outside world [52]. On the contrary, MET takes the ontological problem as fluid, and considers the artefacts as a result of the three factors jointly. Epistemologically, brain, body, and environment not only work together: they are inseparable. Thus, the pot as an action is geared towards the pot as an outcome; the pot as an outcome governs the pot as an action; and the pot as an outcome is that which one assumes is the end point of the action when looking at the pot understood as a trace (see Figure 4, cf. Figure 1, left panel).

**Figure 4.**
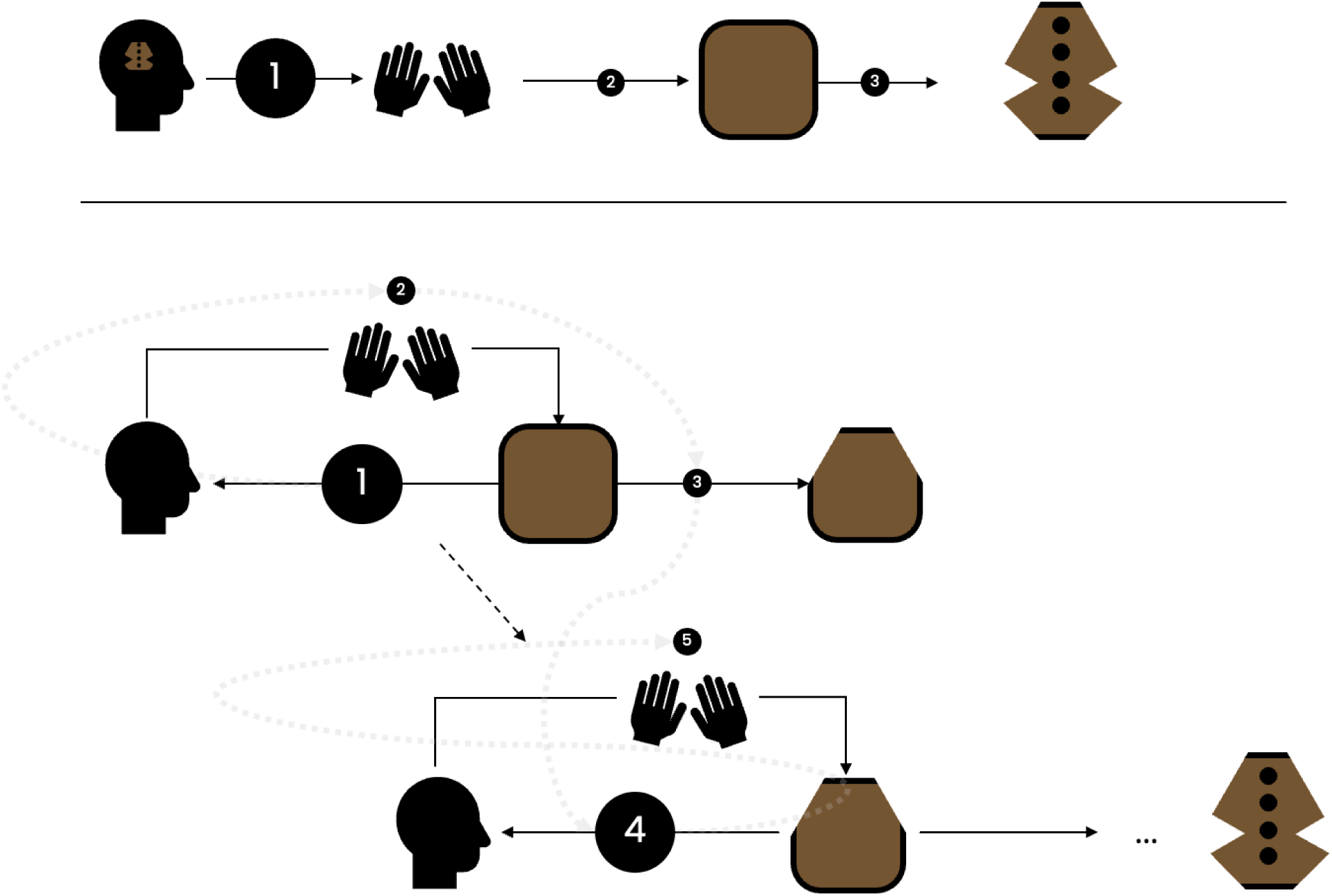
Creative *thinging* under MET. **Top panel:** The “traditional” view assumes that the human maker has a plan “in the head” (1) which is enacted by the activity of her body on the material, (2) turning the material into an outcome, that (3) corresponds to the plan. **Bottom pane**l: MET is the view that the human maker is constrained and affected by the material agent (1) that she engages (2), maybe with a rough idea of (3) producing an outcome. However, in making the material outcome, the outcome approaches what the maker seeks to achieve, and thereby it affords and constrains (4) further embodied interaction, which leads to an outcome (5) that is the result of the joint *thinging* process. Note: The light shaded grey arrows express the nonlinearity of the process.

One way to make sense of the unity of an artefact is to consider artefacts as having a “cognitive life” animated by the “creative *thinging*” of brain, body and environment simultaneously (discussed above, in sec. 1). In creative *thinging* terms, rich pre-planning and pre-representation of outcomes are unnecessary, and the product emerges through a kind of structured give-and-take: the potter continuously nudges the clay, which, in return, continuously nudges the hands of the potter. In this exchange, the embodied interaction is jointly constrained and guided by material signs that both emerge from and structure ongoing activities. The artefact is no longer an “outcome” of a pre-packaged thinking process: it is thinking while feeling and doing. Hence, archaeologists should not seek to interpret the artefact symbolically or functionally, exclusively by trying to infer the hidden intentions of the maker. A more useful way to look at the artefact is to try to dig up how the artefact participated in its own making through the joint process of *thinging*. Thus, the artefact remains also a sign, but a sign of the *thinging* process itself, a way to understand the cognitive unfolding of the artefact-making.

By default, the logic of MET seems incompatible with any form of computation. It is our opinion that this incompatibility can be accommodated, at least partially, by applying AIF, whose tools and logic allow us to analyse phenomena that are relevant to MET. The simulation in this work is the result of this attempt. The key idea was to replicate the process of ‘*thinging*’ through which the human maker leverages and is guided by the material agency of the artefact in-making, while responding to affordances. These affordances are both triggers of complex cultural behavioural patterns, and contemporarily, the negotiation ground that allows the potter to create and innovate. When the materiality offers different levels of uncertainty (for instance, the ludo-engagement with Play-Doh or with Lego, or the carving of a tree branch), the negotiation between the maker and the matter can positively contribute to the creative process. On the other hand, a material which has less uncertainty, that is, one that precisely matches the maker’s engagement (for instance, completing a puzzle or building a cabinet out of a plan), will contribute less to the creation of innovative and creative outcomes.

The between-conditions results indicate how affordances function under varying cognitive and material conditions. Specifically, when operating under less than perfect precision (i.e., a < 1), which aligns with a neurobiological point of view, greater affordance instability (e.g., random or semi-random affordances enabling various interaction patterns) promotes the production of both quite rudimentary and very complex pots compared to conditions where affordances remain stable (e.g., condition **C4**). Notably, there are compelling reasons for the alignment between AIF and MET’s creative *thinging.* In AIF (or “predictive processing”), the brain is fundamentally a ‘lazy’ organ, eager to engage with and use its surrounding reality (see [3]: chapter 8). This occurs because the quality of a predictive model is assessed by accuracy minus complexity, where accuracy refers to the ability to predict the *task-salient* aspects of the sensory information, and complexity indicates the number of parameters in the model. The goal is to learn and use the least complex internal model (with the fewest parameters) that meets our needs [57]. AIF architectures naturally seek the simplest internal models needed for practical use. That is, AIF models will **never** resemble strong instructionist approaches, detailing every step meticulously. Given that perception and action are both driven by the internal prediction machinery simultaneously, actions are shaped by the ultimate goal to support internal-model simplicity. For instance, agents are expected to reach out for the GPS on their phone, rather than wasting precious resources memorising (i.e., learn a complex and expensive internal model) the detailed map of a new city. It is easy to appreciate that this approach strongly favours and actively encourages close cooperation with material resources in the balancing of cognitive resources, the most important of which is attention, or precision weighing.

In our simulation, lowering attention/precision meant focusing less on expected observations, or affordances, during typical pot making trajectories (e.g., being less sensitive to the affordance of “downward-pressability” of the clay at its edge), and more on alternative active patterns (e.g., focusing on the “upward-scraepeability” observation). A lower degree of attention in combination with a “difficult”, or “imprecise” material (for instance, evaluated on a scale going from stable, to semi-stable, and unstable) yielded less aesthetically valuable pieces (given the definition we provided above, see sec. 1). Our results further illustrated a trade-off between precision and affordance instability, revealing a “goldilocks zone” for producing complex artifacts, speaking to the importance of the balancing of cognitive resources in collaboration with material agents. In the semi-unstable affordance condition (**C3**), when precision dropped from a = .8 to a = .7, potter agents shifted from producing more high-value artefacts in **C3** to producing an equal number in both **C3** and **C4**. In our simulation, a = .8 marked the threshold where working with challenging material no longer offered an advantage for creating complex artefacts. Results also showed that in the semi-unexpected affordance condition (**C2**), loss rates were minimised, suggesting that among potters able to make an adequate pot (e.g., pot 2), more were likely to create an apt and highly aesthetic pot (e.g., pot 11).

## 5 LIMITATIONS AND FUTURE DIRECTIONS

This work had to leave out many important factors contributing to the complementarity (or not) of AIF and MET. The first is the ongoing active influence of materiality (‘material agency’) when expert potters work with very well-behaved materials and tools, starting from a clear mental plan. Assuming all goes well on the day, this situation seems to perfectly depict a traditional approach, the one relying upon rich internal representations of the final product, and heavy doses of pre-planning. When a skilled potter starts with a clear idea of the outcome she wants to achieve and deals with compliant materials, detailed management of the necessary potting steps may suffice to achieve a satisfying outcome. We could consider such extreme micro-managing a case of ‘maximal control’ (e.g., precision level a = 1). How should we interpret this situation? Looking at the archaeological past, we find clear examples of vessels whose shapes and decorations were strictly predefined long before their manufacture took place. For instance, the Incan Aryballos (*Aríbalo*), some of the most frequent vessels in the Incan state, were extremely standardised in form and decoration [58–61]. Yet, despite the strictly predetermined shape, they were produced adopting various techniques and material components in the different regions of the Incan empire. For instance, in the Puna region (Argentina, Bolivia, and Chile), Incan-style vessels show the addition of pelites in the clay, a distinctive feature of the Yavi tradition [62]. This is because, to facilitate the production and, ultimately, the spreading of the Aryballos vessels, their manufacture depended upon pre-existing potter traditions [63]. Consequently, potters had to make continuous adjustments to their usual repertoire to fulfil the requirements of the imperial vases. That is, even in the presence of pre-designed templates and standardised pottery vessels demanded by imperial authorities, those artefacts show the continuous negotiation between the potter, the material, and the artefact itself. That is, also in cases of ‘maximal control’, the bedrock picture remains the same: despite the surface appearances, they are the result of the *thinging* process.

The explanation is both exciting and enlightening. As discussed earlier (sec. 1 and 4), the human potter learns to control the clay through an inner model that optimally balances accuracy and complexity. This model enables her to get the job done using the least complex set of internal resources, considering the conditions encountered during learning. Consequently, it focuses on maximising the capabilities of tools and the inherent dynamics of materials. To make the concept clearer, we can exemplify using the passive dynamics of the human body when controlling walking, running, and even climbing. By making the most of the shape and connectivity of bones, muscles, and tendons, the task of controlling stable powered locomotion is greatly simplified [64–66]. Neural control systems co-evolved with human bodies, allowing each side to contribute in ways that together enable fluid, energetically-efficient locomotion. For an even simpler example, consider the ease with which a simple ‘slinky spring’ (stairwalker) uses gravity and its intrinsic dynamics to power an elegant descent – one that would otherwise require very large amounts of computation to pre-program! Albeit at a different timescale, it is in the same way that the potter’s brain learns how to make the most of a material (the clay) that has been selected as a good partner for the task of (roughly speaking) vessel production. The internal models that then arise will be a very far cry from the kind of pre-imagined micro-management control techniques rightly dismissed by MET. Instead, the control model that emerges will make the most of all the rich dynamics of the clay, and this will be the case even if (as in the maximal-control case above) they are simply engaging with the clay to bring about a richly pre-imagined outcome. Notice also that even in bringing about a richly pre-imagined outcome, the potter will not pre-think every single step along the way. Instead, they will have learnt a set of cheap, effective recipes for engaging the medium such that the next step becomes apparent (is computed) only after an earlier step is taken.

Now, let’s consider a different set of cases, when things don’t quite go according to plan. It is when things go sideways that true expertise manifests. In such cases, prediction error signals arise and highlight the mismatch between expectations and sensory inputs (what is sensed and what was, at that moment in time, expected to be sensed). These signals are processed until remedial actions can be calculated, restoring alignment. This is where AIF excels: while it supports fast, efficient (‘habitual’) responses, it reveals that such actions are simply one ‘setting’ in an overall cognitive economy that constantly self-organises around prediction error signals. If prediction error signals form and dissolve just as expected, behaviour is fluid and often relies on very low-cost (as far as internal processing is concerned) strategies. Truly unexpected errors, however, engage deeper (and deeper) processing and can draw upon additional knowledge and skill-sets, enabling the expert to try out new moves and potentially discover new solutions. Thus, low-cost solutions that make maximal use of body and world co-exist with a range of potential ‘high-cost’ strategies that involve stepping back and thinking more reflectively, and all points in between. That said, the expert potter’s brain will not usually need to push prediction errors very far up the line to recruit an apt response when things start to drift out of line. Most importantly, all responses (from the highly reflective to the seemingly automatic) are curated in the same way, by self-organising around prediction error signals during embodied interactions with the world (For a fuller discussion of this, see: [67]; [3]).

What, finally, does all this suggest about ‘creativity’ itself? Creativity, we conjecture, is often curated by deliberately ‘holding loose’ some things that might otherwise be more highly controlled [54]. Here, AIF’s ‘precision-weighting’ dimensions again provide a useful framework. Imagine a potter capable of engaging a specific material (a highly standardised clay) with maximum control. To enable more creative explorations, the potter might lower the precision assigned to their own expectations, and they might do so at variable levels. They might lower the precision on the overall outcome, or on any of the many recipes for engagement that arise along the way. They might also alter the materiality itself, deliberately deploying some new mixture or new tools, simply to see what happens. All these ploys fall into place as ways of allowing more voice to different aspects of materiality or practice. For instance, potters from the Raku tradition in Japan allow for cracks during the firing process to define the final ‘look’ of the vessels, so that a certain degree of unpredictability is integrated in the design of the artefact [68,69]:307;[68,69]. Raku pottery consisted of small hand-made and lead-glazed tea cups fired in small cylindrical kilns [70], and is most famously known for its unique (traditionally low-temperature) firing technique. During this process, potters are constantly monitoring the fuel intake, as well as the flames produced, and after the firing is thought to be enough, the pots are quickly removed from the kiln, displaying a spectacular glow.

At this moment, the pots start taking their final look, and there is great variation in the results of each firing [68]. Crazing and other faults produced at this point are considered part of the aesthetics of ‘imperfection’, and highly valued. Therefore, to a degree, the firing in itself is controlled, sometimes to incredible extents, but the glazing of the pot’s surface is let loose, i.e. left to the whims of the materials’ touch with the environment. As a result, unique designs are created.

Collective human endeavour adds yet more dimensions that warrant future examination. Groups of human agents seem to act to minimise a kind of ‘collective prediction error’ thanks to iterated interactions within a socially structured space – as when a ‘pottery collective’ slowly develops and refines a new style of pottery. Here too, the same basic toolkit should apply (for some preliminary explorations, see [28]). In future work, we plan to use the simple platform described in the present paper to explore all these (and other) cases even further, revealing AIF and MET as complementary perspectives on the ways materiality, embodied action, and neuronal processing conspire to enable adaptive success.

### Software note

The implementation of variational message passing is done with the open-source code commonly used in active inference simulation studies, accessible as a MATLAB routine (spm_MDP_VB_X.m) in the SPM12 toolbox. The routine takes in as inputs a structure of prespecified initial state (D), likelihood (A) and transition matrices (B), as well as prior preference (C) and set policies (V) and returns a sequence of belief updates and actions that minimise variational and expected free energy.

## Footnotes

1 Claudio Tennie’s ‘test ìsland’ studies are a beautiful example of how righ-preplanning or sophisticated cognition is not necessary in the making of artefacts that are often interpreted as representation of complex cognitive and social abilities, and that even in humans some environmental conditons (e.g., social and technical availability of models) might be necessary for producing sophisticated artefacts (see, e.g., [6–11]).

2 While we are aware that, in reality, such a cultural process invests every aspect of the production of an artefact - from the tools that are used to the working methodology, to the selection of the raw material itself, such a complexity had to be necessarily simplified in the present work. For satisfying the curiosity of the reader, we might suggest to look at, e.g.,[30]; [31] [32]; [33].

3 This sequence provides flexibility compared to the traditional sequence of pottery manufacture. In traditional pottery-making, the decoration is decidedly kept as a final process, occurring only after vessels have been fully formed and partly left to dry. While there are technical and physical aspects to conceptualising pottery manufacture through such sequences, the idea was not to restrict the simulated potter to a rigid sequence of production and enable possibilities informed by the lump of clay for making adjustments as the manufacturing process unfolds.

4 Indirectly, it might also suggest that the level of the precision imposed by the potter or human agent, in terms of expertise, might affect the final result.

